# Characteristics of JN.1-derived SARS-CoV-2 subvariants SLip, FLiRT, and KP.2 in neutralization escape, infectivity and membrane fusion

**DOI:** 10.1101/2024.05.20.595020

**Authors:** Pei Li, Julia N. Faraone, Cheng Chih Hsu, Michelle Chamblee, Yi-Min Zheng, Claire Carlin, Joseph S. Bednash, Jeffrey C. Horowitz, Rama K. Mallampalli, Linda J. Saif, Eugene M. Oltz, Daniel Jones, Jianrong Li, Richard J. Gumina, Kai Xu, Shan-Lu Liu

## Abstract

SARS-CoV-2 variants derived from the immune evasive JN.1 are on the rise worldwide. Here, we investigated JN.1-derived subvariants SLip, FLiRT, and KP.2 for their ability to be neutralized by antibodies in bivalent-vaccinated human sera, XBB.1.5 monovalent-vaccinated hamster sera, sera from people infected during the BA.2.86/JN.1 wave, and class III monoclonal antibody (Mab) S309. We found that compared to parental JN.1, SLip and KP.2, and especially FLiRT, exhibit increased resistance to COVID-19 bivalent-vaccinated human sera and BA.2.86/JN.1-wave convalescent sera. Interestingly, antibodies in XBB.1.5 monovalent vaccinated hamster sera robustly neutralized FLiRT and KP.2 but had reduced efficiency for SLip. These JN.1 subvariants were resistant to neutralization by Mab S309. In addition, we investigated aspects of spike protein biology including infectivity, cell-cell fusion and processing, and found that these subvariants, especially SLip, had a decreased infectivity and membrane fusion relative to JN.1, correlating with decreased spike processing. Homology modeling revealed that L455S and F456L mutations in SLip reduced local hydrophobicity in the spike and hence its binding to ACE2. In contrast, the additional R346T mutation in FLiRT and KP.2 strengthened conformational support of the receptor-binding motif, thus counteracting the effects of L455S and F456L. These three mutations, alongside D339H, which is present in all JN.1 sublineages, alter the epitopes targeted by therapeutic Mabs, including class I and class III S309, explaining their reduced sensitivity to neutralization by sera and S309. Together, our findings provide insight into neutralization resistance of newly emerged JN.1 subvariants and suggest that future vaccine formulations should consider JN.1 spike as immunogen, although the current XBB.1.5 monovalent vaccine could still offer adequate protection.

## INTRODUCTION

Tracking the ongoing evolution of SARS-CoV-2 and its impacts on spike protein biology, particularly sensitivity to neutralizing antibodies, is critical as the pandemic continues. The pandemic underwent a turning point in late summer 2023 with the emergence of BA.2.86, a variant characterized by over 30 spike protein mutations relative to then dominate variant XBB.1.5^1^. Fortunately, despite its myriad mutations, BA.2.86 did not exhibit increased immune evasion, but was better neutralized by antibodies in convalescent and vaccinated sera relative to XBB-lineage variants^2–11^. However, mounting concern has arisen with the subsequent variants that have evolved from BA.2.86. This includes JN.1, which emerged in late 2023 and is characterized by the single spike mutation L455S relative to BA.2.86^1^. This single mutation launched JN.1 to dominance worldwide from late 2023 through May 2024^12^. L455S contributes to the lower affinity of JN.1 for human ACE2 but enhances its immune evasion to neutralizing antibodies and viral transmission^9,13–18^.

Since JN.1’s emergence, a series of variants that possess mutations at key sites in spike have been identified, including L455, F456, and R346 (**Figure 1A**). Initially, the so-called FLip variants emerged, possessing L455F and F456L mutations in the backbone of XBB.1.5, hence the name “FLip” ^5,10^. These sites have continued to be hotspots, with a strain called “SLip” having emerged, which has the JN.1 spike protein with the F456L mutation - the “S” referring to the L455S mutation that characterizes JN.1. More recently, we have seen the emergence of the FLiRT variant, which harbors an additional R346T mutation in the backbone of SLip. Another variant, called KP.2, contains both R346T in S1 as well as an V1140L mutation in S2. JN.1 is currently waning in dominance around the world, becoming quickly supplanted in circulation by KP.2 and other JN.1 derived variants^12,19^ (**Figure 1B**).

**Figure 1:**
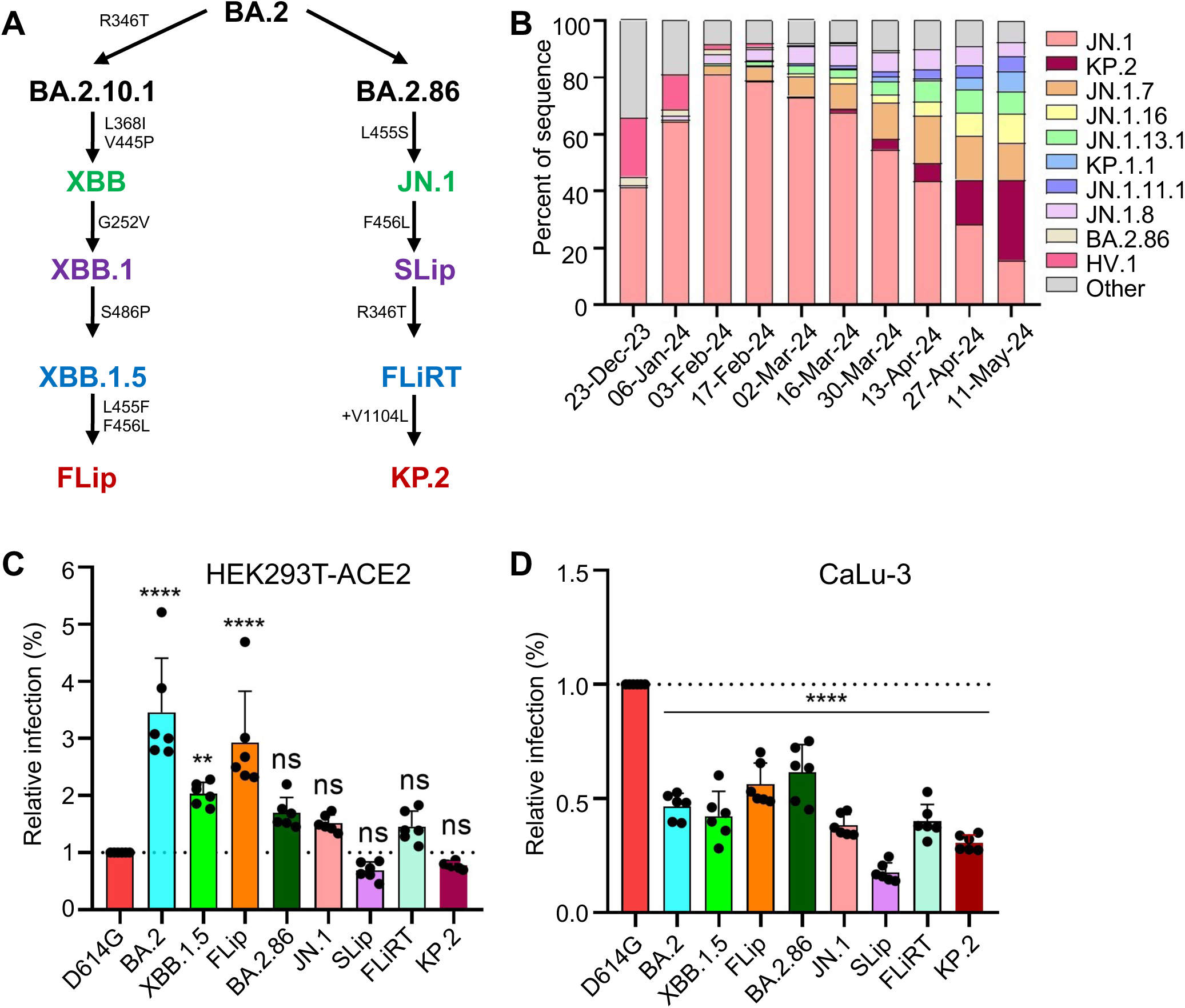
Genetic relationship, distribution and infectivity of JN.1-derived subvariants. **(A)** Schematic depicting spike protein mutations that characterize JN.1 and its subvariants. Related XBB.1.5 variants including FLiP are included. **(B)** Variant proportions over time in circulation in the United States (December 2023 – May 2024). Data was downloaded from the Centers of Disease Control website and replotted. **(C-D)** Infectivity in 293T-ACE2 and CaLu-3 cells. Pseudotyped lentiviruses bearing the spike of interest were used to determine entry into **(C)** 293T-ACE2 and **(D)** CaLu-3 cells. Relative luminescence readouts were normalized to D614G (D614G = 1.0) for plotting. Bars in (C) and (D) represent means ± standard deviation from six individual measurements of viral infection of different doses (n = 6). ** p < 0.01; ****p < 0.0001; ns p > 0.05.

It has been shown that JN.1 can be neutralized by XBB.1.5-monovalent vaccinated sera, albeit with a reduced efficiency^9,16,17,20^. However, it is currently unclear whether additional spike mutations gained in the JN.1 lineage subvariants will affect the efficacy of this COVID-19 vaccine formulation, especially with the approach of upcoming fall and winter seasons. In this study, we investigated the ability of SLip, FLiRT, and KP.2, in parallel with their parental JN.1, to be neutralized by sera from: (i) individuals vaccinated with at least 2 doses of monovalent ancestral spike (wildtype, WT) mRNA vaccine with 1 dose of bivalent WT+BA.4/5 booster, (ii) hamsters vaccinated with 2 doses of XBB.1.5 monovalent recombinant mumps vaccine, and (iii) individuals infected during the BA.2.86/JN.1-wave of infection in Columbus, OH. These analyses were conducted alongside the ancestral D614G and are supplemented with antigenic cartography analyses. We also characterized the entry and fusogenicity of these variants in 293T-ACE2 and CaLu-3 cells, as well as the spike processing and expression on plasma membranes. Critically, our investigation focuses on the comparison between XBB.1.5 and JN.1 as immunogens against these JN.1-lineage variants and determines whether any dramatic changes in spike protein biology have occurred, as well as their impact on neutralization escape and viral infectivity.

## RESULTS

### Impacts of JN.1-derived variants on viral entry and infectivity in 293T-ACE2 and CaLu-3 cells

We first investigated the efficiency in which pseudotyped lentiviruses bearing SARS-CoV-2 spikes of interest can enter 293T cells overexpressing human ACE2 (293T-ACE2) and human lung epithelial cell line CaLu-3. As reported previously by our group, earlier Omicron variants, including BA.2 and XBB.1.5, exhibited higher infectivity relative to the ancestral D614G variant in 293T-ACE2 cells^21^; however, infectivity decreased modestly for BA.2.86 and JN.1 (**Figure 1C**)^16^. Here we found that while FLiRT exhibited a similar infectivity to JN.1 in this cell line, KP.2 (p > 0.05), and especially SLip (p < 0.05), demonstrated modest reductions compared to JN.1 (**Figure 1C**). As we demonstrated previously, Omicron variants maintain a markedly reduced infectivity in CaLu-3 cells relative to D614G, with a notable increase for BA.2.86^21,22^. However, we observed here that JN.1, SLip, FLiRT and KP.2 exhibited decreased infectivity in CaLu-3 cells compared to BA.2.86 (p < 0.0001 for all), with SLip exhibiting the lowest infectivity of the group, ∼2.2-fold decrease compared to JN.1 (p < 0.001) (**Figure 1D**). Overall, the recently emerged FLiRT and KP.2 subvariants, especially SLip, exhibit decreased infectivity in CaLu3 cells compared to earlier variants BA.2 and XBB.1.5, as well as their parental BA.2.86.

### Increased resistance of SLip, FLiRT and KP.2 to COVID-19 bivalent vaccinated human sera by newer variants compared to parental JN.1

To test the extent of escape from neutralization by selected variants, we first used sera were from a cohort of healthcare workers (HCWs) at The Ohio State University Wexner Medical Center, who had received at least 2 doses of monovalent mRNA vaccine (WT) plus at least 1 dose of bivalent vaccine containing both WT and BA.4/5 spikes (n=10) (**Figures 2A-B**). Neutralization was measured using pseudotyped lentiviruses mixed with serial dilutions of sera and infected 293T-ACE2 cells to determine neutralization titers at 50% (NT_50_) for each variant spike. As shown for all previous Omicron variants^5,16,23^, JN.1 had markedly lower titers relative to the ancestral D614G, with a 53.3-fold decrease (p < 0.0001). SLip, FLiRT, and KP.2 also exhibited dramatically decreased titers, with NT_50_ 56.3-fold (p < 0.0001), 86.4-fold (p < 0.0001), and 76.7-fold (p < 0.0001) lower than D614G, respectively. Notably, whereas SLip showed a similar titer to JN.1, FLiRT and KP.2 had more dramatic decreases in titer, with 1.62-fold (p > 0.05) and 1.43-fold (p > 0.05) lower than JN.1, respectively (**Figures 2A-B**). Overall, FLiRT and KP.2 exhibit increased escape from neutralizing antibodies in bivalent vaccinated sera compared to parental variant JN.1 and related variant SLip.

**Figure 2:**
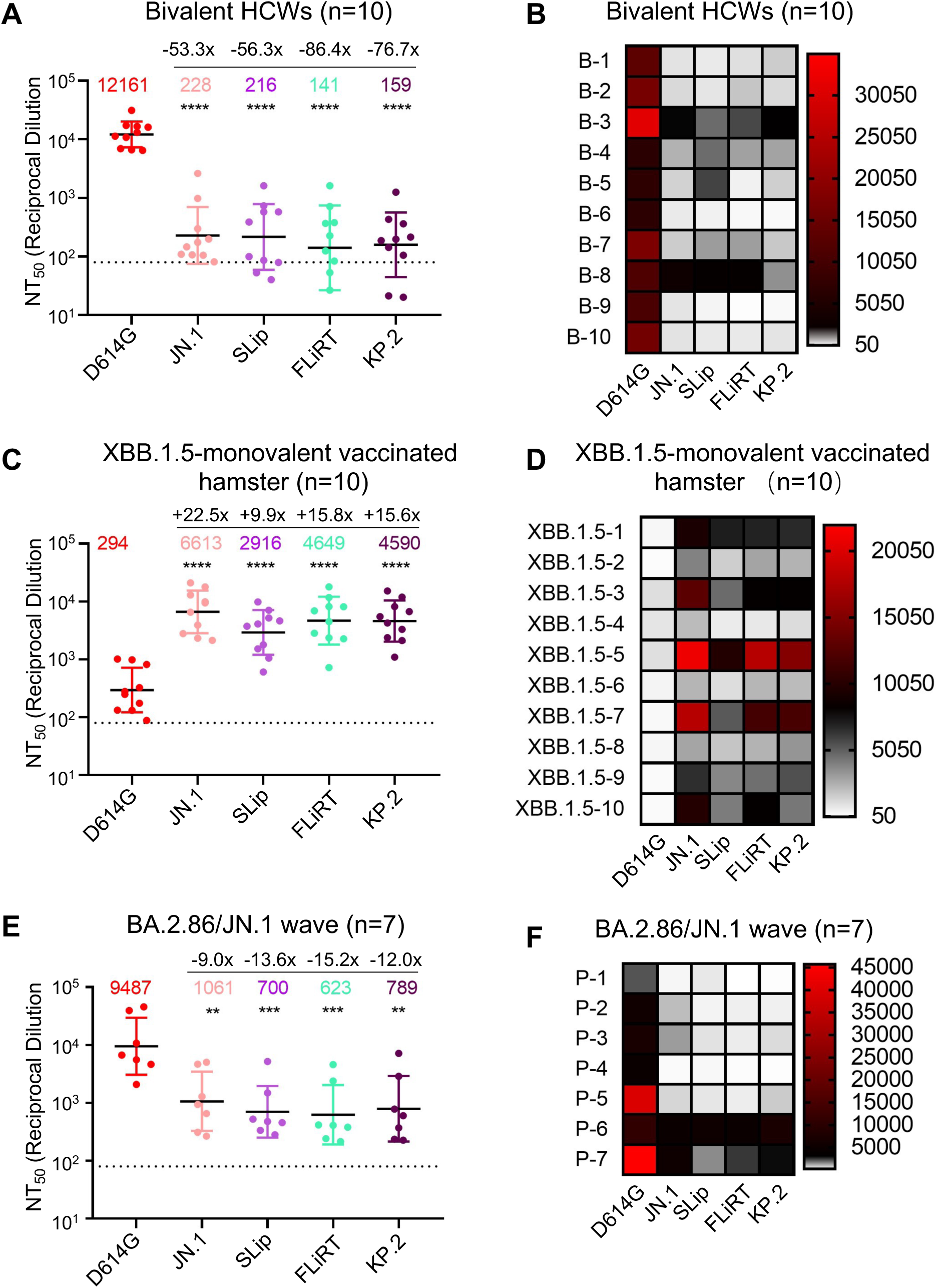
Neutralization of SLip, FLiRT, and KP.2 by bivalent vaccinated healthcare workers, XBB.1.5-vaccinated hamsters, and BA.2.86/JN.1 convalescent individuals. Pseudotyped lentivirus was used to perform neutralizing antibody assays with **(A-B)** sera from HCWs that received at least 2 doses of monovalent vaccine and 1 dose of bivalent booster ( n= 10), **(C-D)** golden Syrian hamsters inoculated twice with recombinant mumps virus carrying XBB.1.5 spike (n = 10), **(E-F)** people infected during the BA.2.86/JN.1-wave in Columbus, OH, and **(A, C, and E)**. Plots depicting the geometric mean of neutralizing antibody titers at 50% (NT_50_) are shown, with fold changes relative to D614G displayed at the very top. **(B, D, and F)** Heatmaps depicting the NT_50_ values for each cohort by individual sera sample. **p < 0.01; ***p < 0.001; ****p < 0.0001.

### Antibodies in XBB.1.5 monovalent vaccinated hamster sera robustly neutralize FLiRT and KP.2, with reduced efficiency for SLip

We next measured neutralization using sera from golden Syrian hamsters vaccinated twice with XBB.1.5 monovalent recombinant mumps vaccine (n=10) (**Figures 2C-D**). As demonstrated previously^16^, the hamster sera had robust neutralizing antibody titers against some of the latest Omicron subvariants, including BA.2.86, as compared to the ancestral D614G. Neutralization titers for JN.1 were also increased, with a calculated NT_50_ value of 6,613, which was 22.5-fold high than D614G (p < 0.0001). The NT_50_ values for SLip, FLiRT, and KP.2 showed 9.9-fold (p < 0.0001), 15.8-fold (p <0.0001), and 15.6-fold (p < 0.0001) increases compared to D614G, respectively, corresponding to a 2.3-fold, 1.4-fold and 1.4-fold decrease, respectively, relative to JN.1 (p > 0.05 for each) (**Figures 2C-D**). The overall high titer of XBB.1.5 monovalent vaccinated hamster sera against these new JN.1-derived variants was in sharp contrast to the generally low antibody titer exhibited by the bivalent vaccinated group (**Figures 2A-D**), but the downward trends of each variant were similar, except for SLip. The relatively strong neutralization escape of SLip from XBB.1.5 monovalent vaccine, as compared to FLiRT and KP.2, was likely due to a specific amino acid R346T change in the receptor-binding domain of XBB.1.5, FLiRT and KP.2 spikes, but not in the SLip spike (see *Discussion*). Nonetheless, sera from XBB.1.5-vaccinated hamsters can effectively neutralize JN.1 and JN.1-derived subvariants.

### SLip, KP.2, and especially FLiRT, exhibit decreased sensitivity to neutralization by BA.2.86/JN.1-wave convalescent sera compared to parental JN.1

To examine variant neutralization by antibodies induced during a natural infection, we employed sera from individuals who tested positive for COVID during the BA.2.86/JN.1 wave of infection during November 2023 and February 2024 in Columbus, Ohio (n=7) (**Figures 2E-F**). Four were Columbus first-responders and their household contacts (n=4, P1 to P4) and three were ICU COVID-19 patients admitted to the OSU Medical Center (n=3, P5 to P7). All patients had received different doses of mRNA vaccine, with samples being collected between 34-892 days following the last vaccination (**Table S1**). As we have shown previously^16^, neutralization titers were detectable, albeit modest, and especially for P1-P4 against JN.1, with about 9-fold reduction relative to D614G (p < 0.01). Titers against SLip, FLiRT, and KP.2 were further decreased, with reductions of 13.6-fold (p < 0.001), 15.2-fold (p < 0.001), and 12-fold (p < 0.01) relative to D614G, respectively. Similar to the bivalent cohort, the FLiRT variant exhibited the biggest drop in titers compared to JN.1 (1.70-fold decrease), though the difference was not statistically significant (p > 0.05) (**Figures 2E-F**). Samples P5, P6, and P7 were collected from individuals admitted to the ICU at the Ohio State University Wexner Medical Center (**Figure 2F, Table S1**). Notably, two of these patients, P6 and P7, exhibited higher titers against the JN.1-lineage variants. P6 is a 77-year-old male who received one dose of the Moderna monovalent vaccine and one dose of the Pfizer bivalent vaccine, with the sample taken 434 days after his last vaccination. P7 is a 46-year-old female who received three doses of the Moderna monovalent vaccine and one dose of the Moderna bivalent vaccine, with her sample collected 334 days after her last vaccination. P5 is a 49-year-old male ICU patient and had only received two doses of the Moderna monovalent vaccine; his sample was taken 892 days after his last vaccination, with neutralizing antibody titers against JN.1, SLip, FLiRT, and KP.2 being the lowest among the three ICU patients. Overall, sera of BA.2.86/JN.1 convalescent individuals effectively neutralized the latest JN.1-lineage subvariants SLip, FLiRT and KP.2, but with somewhat reduced efficiency for FLiRT.

### Class III monoclonal antibody (mAb) S309 does not neutralize SLip or FLiRT

Another critical strategy for pandemic control measures is the use of therapeutic monoclonal antibodies, which was demonstrated during COVID-19 pandemic, especially between 2020-2021^24^. However, because of their binding being limited to a single epitope on the SARS-CoV-2 spike, single mutations can easily disrupt their efficacy, making most of the developed mAbs completely ineffective^25–27^. Notably, we and others have previously shown that binding of class III mAb S309 is largely maintained against Omicron subvariants, apart from BA.2.75.2, CA.3, CH.1.1, BA.2.86, and JN.1^5,16,23,28^. This trend appeared to continue, as both SLip and FLiRT exhibited a complete escape of neutralization by S309 (**Figures 3A-B**). We did not perform this experiment for KP.2, which harbors the conserved D339H mutation critical for S309 resistance (see *Discussion*).

**Figure 3:**
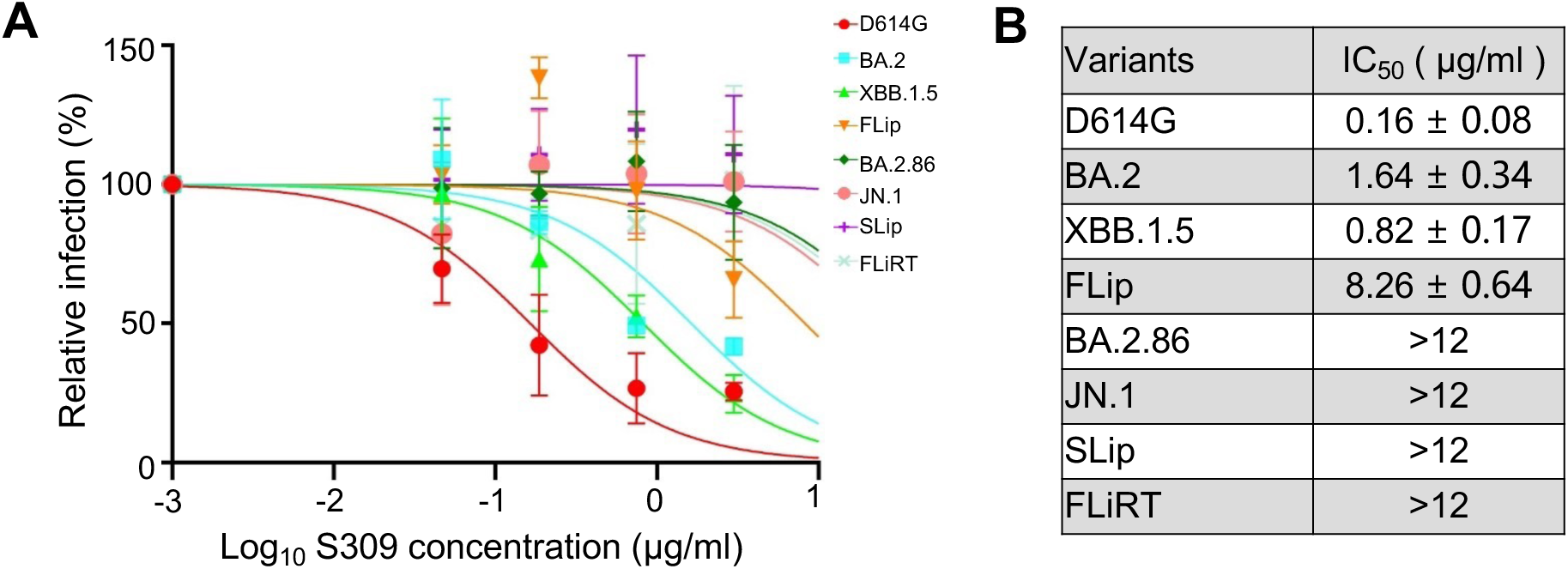
Neutralization of SLip, FLiRT, and KP.2 by class III mAb S309. Neutralization was performed using lentiviral pseudotypes carrying each of the indicated spike proteins of the JN.1 subvariants to assess the effectiveness of mAb S309. **(A)** Neutralization curve for each of the variants for S309 and **(B)** the calculated IC_50_ values.

### Antigenic cartography analysis demonstrates decreased antigenic distances of SLip, FLiRT and KP.2 in XBB.1.5 monovalent-vaccinated and BA.2.86/JN.1-infection groups

To further elucidate the relationships between these variants, we conducted antigenic cartography analysis (**Figure 4**). Briefly, this analysis transforms neutralization titers based on relative differences between titers for each variant (circles) and each serum sample (squares) displayed as antigenic units (AU). As would be expected, JN.1 subvariants are antigenically distinct from the ancestral D614G in the bivalent cohort^16^, with FLiRT being the most distinctive compared to SLip, KP.2 and their parental JN.1 (**Figure 4A**). In the XBB.1.5 monovalent vaccinated hamster group, distances between D614G and the JN.1 variants are markedly reduced, from ∼6 AU in the bivalent cohort down to ∼4-5 AU. The variants are also clustered more closely to each other, with SLip being slightly further away from FLiRT and KP.2 (**Figure 4B**). A similar phenomenon was observed in the BA.2.86/JN.1 wave cohort, with overall shorter antigenic distances (∼3-4 AU) compared to the bivalent and XBB.1.5 monovalent cohorts. Again, JN.1-derived subvariants largely cluster together, with parental JN.1 being relatively distant from SLiP, FLiRT, and KP.2 (**Figure 4C**).

**Figure 4:**
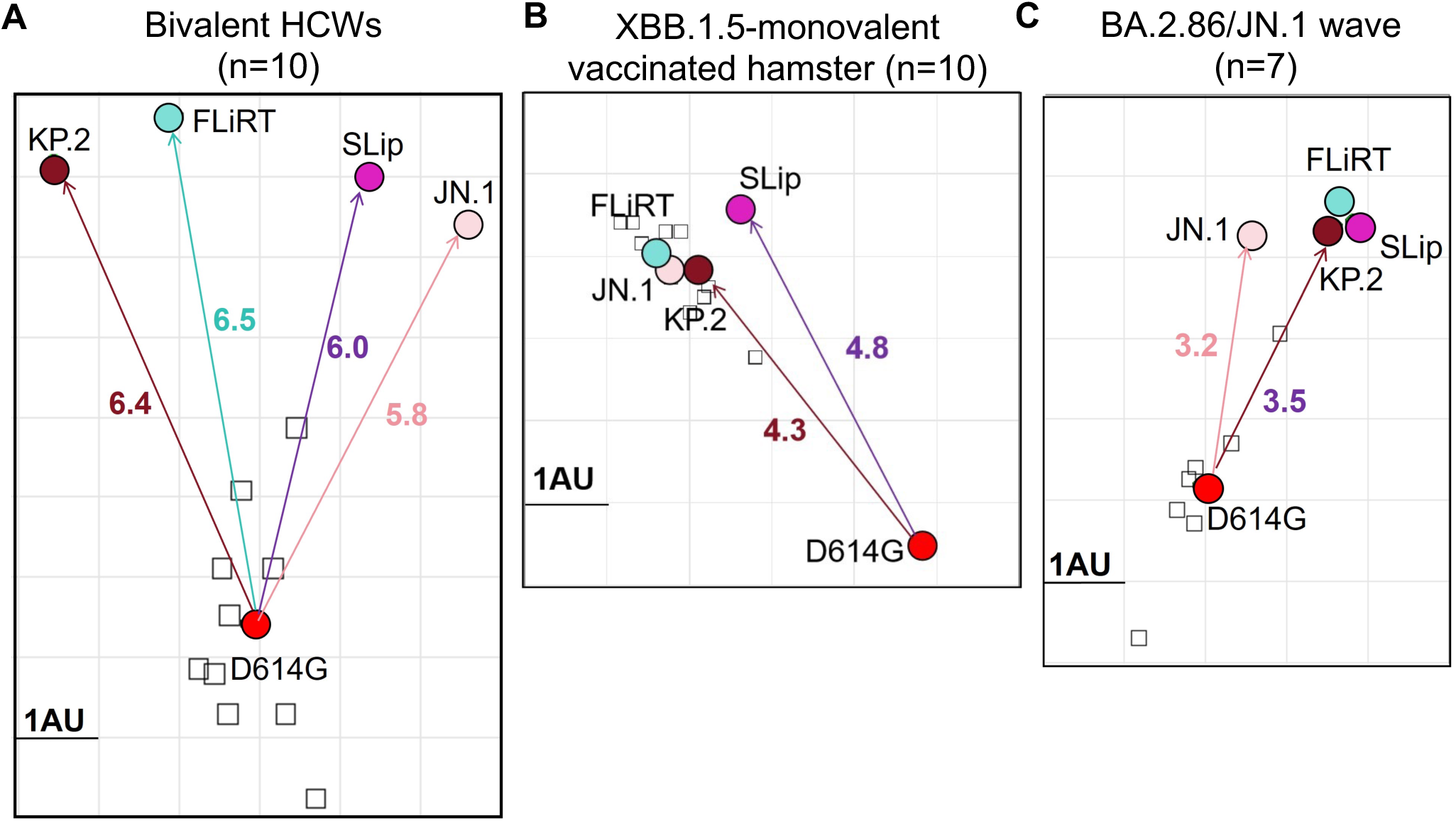
Antigenic mapping of neutralization titers for bivalent-vaccinated, XBB.1.5 monovalent-vaccinated, and BA.2.86/JN.1-wave-infected cohorts. Antigenic cartography analysis was conducted using the Racmacs program to create antigenic distance maps for the neutralization titers in the **(A)** bivalent HCW, **(B)** XBB.1.5-monovalent vaccinated hamsters, and **(C)** the BA.2.86/JN.1 convalescent cohorts. Colored circles represent the different spike antigens, small squares represent individual sera samples. One antigenic distance unit (AU = 1) is represented by one side of the grid squares. 1 AU is equivalent to about 2-fold differences in overall neutralization titers.

### SLip, FLiRT, and KP.2 spikes exhibit modestly decreased fusogenicity, surface expression and processing relative to JN.1

Our previous studies revealed notable changes in spike biology of many Omicron variants, including their membrane fusogenicity and processing. Here we characterized the ability of spikes from new JN.1-derived subvariants to trigger fusion between cell membranes (**Figures 5A-D**), their expression on cell plasma membranes (**Figures 5E-F**), as well as their processing into S1/S2 subunits by furin in virus producer cells (**Figure 5G**). Similar to other Omicron variants, JN.1 exhibited markedly reduced cell-cell fusion activity relative to D614G^16^. This downtrend was maintained for FLiRT and KP.2, and more so for SLip, in both 293T-ACE2 and CaLu-3 cells (p < 0.0001 compared to D614G). The level of cell-cell fusion activity for these three new subvariants appeared to be lower than parental JN.1 (p < 0.01). We also investigated expression of spikes on the surface of 293T cells, which were used to produce pseudotyped vectors. We found that JN.1-derived subvariants exhibited a 2∼3-fold decrease in expression compared to ancestral D614G (p < 0.0001), with FLiRT being significantly lower than the parental JN.1 (p < 0.0001) (**Figures 5E-F**). We probed lysates of the 293T cells for S2 subunits of spike to determine the extent of furin cleavage efficiency by quantifying the ratio of S2/S, and we observed that the processing efficiency for SLip, FLiRT, and KP.2 spikes was modestly decreased compared to JN.1 (**Figure 5G**). Overall, the decreases in cell-cell fusion and spike processing are consistent with their attenuated infectivity in 293T-ACE2 and CaLu-3 cells (**Figures 1C-D**).

**Figure 5:**
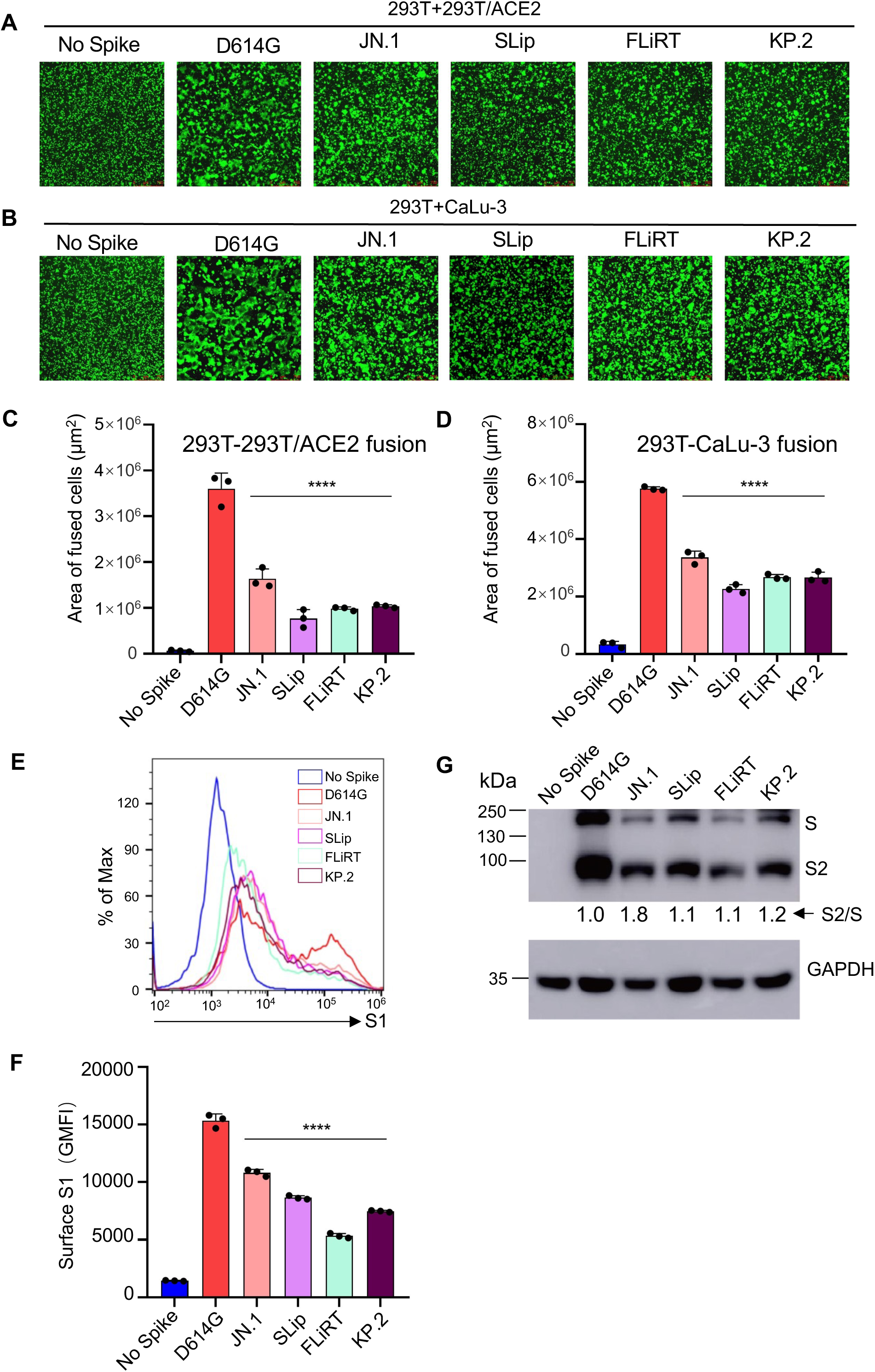
Cell-cell fusion, surface expression and processing of SLip, FLiRT, and KP.2 spikes. **(A-B)** Representative images of fused cells with 293T cells transfected to produce spike plus GFP and co-cultured with **(A)** 293T-ACE2 cells or **(B)** CaLu-3 cells. Images were taken 4 hours (CaLu-3) and 6.5 hours (293T-ACE2) after co-culturing. **(C-D)** Plots of average area of fused cells for each spike for 3 total replicates (n = 3) for **(C)** 293T-ACE2 and **(D)** CaLu-3 cells. **(E-F)** Surface expression of spike on 293T cells used to produce pseudotyped lentiviruses was determined using anti-S1 antibody by flow cytometry. **(E)** Representative histogram depicting relative S1 signal for each variant and **(F)** a plot of the geometric mean fluorescence values for 3 replicates (n =3). **(G)** 293T cells used to produce pseudotyped lentivirus were lysed and used for western blotting to probe for full length and S2 subunits of spike and GAPDH (loading control). Relative differences between band intensities were determined using NIH Image J and normalized to D614G (D614G = 1.0). ****p < 0.0001.

### Structural modeling of mutations in SLip, FLiRT, and KP.2 spikes

To better understand the impact of spike mutations on these new variants, we performed homology modeling to investigate alterations in receptor engagement, spike conformational stability, and antibody interactions. Residue R346 in the receptor binding motif (RBM) can form both a hydrogen bond and a salt bridge with residue D450, which is present in the parental BA.2.86 lineage (N450D). This interaction pulls residue D450 and its loop away from RBM and potentially disturbs the process of receptor engagement. The R346T mutation abolishes this interaction, releasing tension on residue D450 and the RBM, thus potentially enhancing ACE2 binding affinity (**Figure 6A**). Conversely, residues L455 and F456, which are centrally located within the RBM, are encased in a hydrophobic cage formed by Y421, Y453, Y473, and Y489. This hydrophobic core is crucial for ACE2 binding. Mutations such as F456L and L455S found in strains JN.1 and SLip can reduce the local hydrophobicity of the RBM, diminishing interactions with ACE2 residues T27, K31, D30, and H34 and thus potentially decreasing viral affinity for ACE2 (**Figure 6B**). Additionally, structural analysis indicates that residue V1104 is situated in a hydrophobic core, together with P1090, F1095 and I1115 on the spike stem region. The V1104L mutation fills a cavity and improves the local hydrophobic interaction, potentially stabilizing the prefusion spike conformation, which would reduce the efficiency of spike protein transition to a postfusion conformation (**Figure 6C**). Residues D339 and R346 lie within the epitope region of class III antibodies, including S309 (**Figure 6D**). Mutations at these positions are present in the BA.2.86 lineage and subsequent lineages FLiRT and KP.2, which could enhance viral evasion from antibody neutralization. Lastly, residues F456 and L455 on the RBM are frequently targeted by class I RBD neutralizing antibodies, such as CC12.1 (**Figure 6E)**. Therefore, the L455S and F456L mutations, which involve changes in size and chemical properties, can effectively enable viral evasion from humoral immunity established by prior infection and/or vaccination.

**Figure 6:**
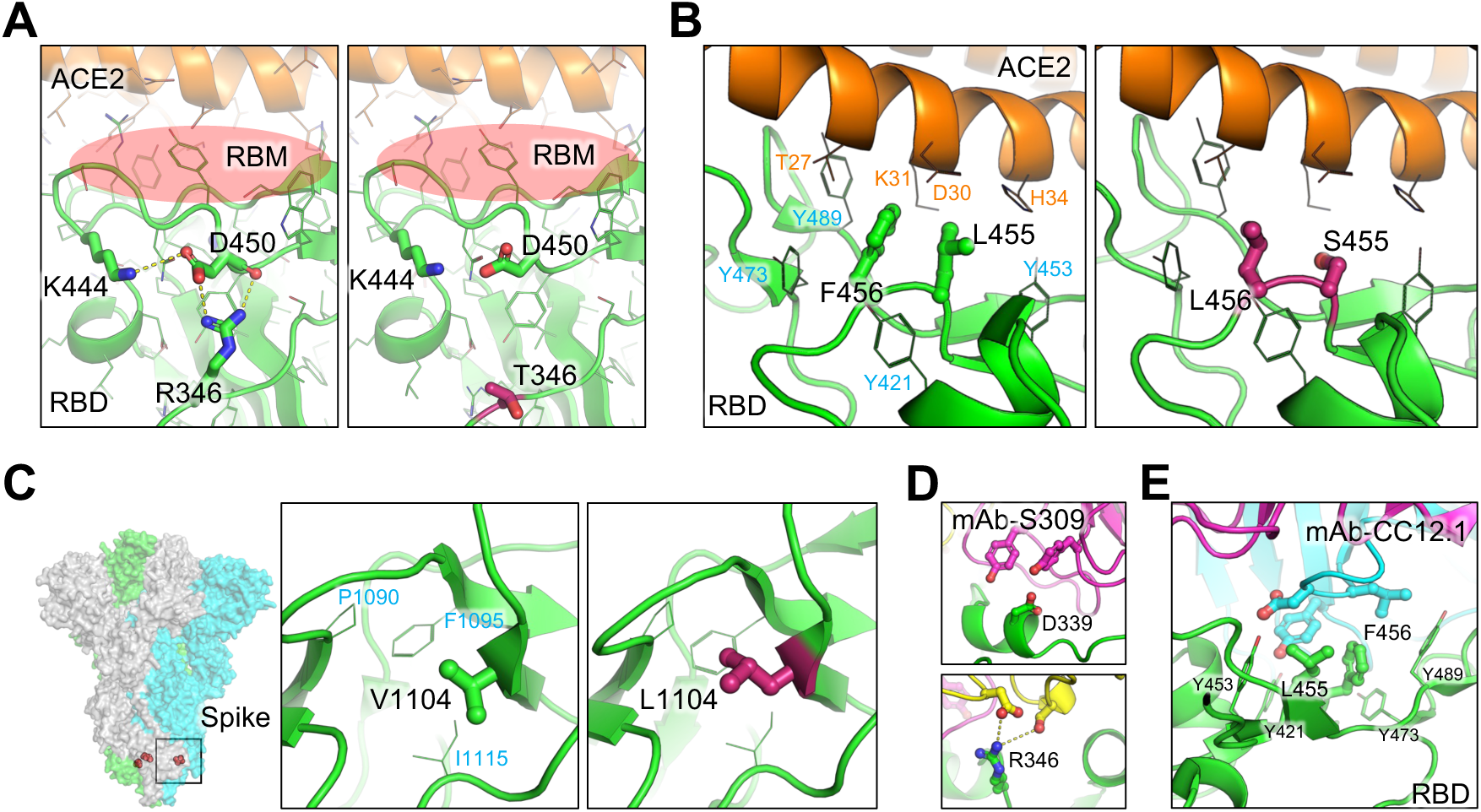
Structural modeling of ACE2 binding, conformation stability, and antibody evasion by mutations on SLip, FLiRT, and KP.2 spikes. **(A)** R346T enhances ACE2 binding by abolishing a salt bridge to D450 and releasing the tension on RBM. Contacting residues are shown as sticks. Hydrogen bonds and salt bridge are shown as yellow dots. **(B)** F456L and L455S reduce the local hydrophobicity on RBM, therefore potentially decreasing the spike affinity for ACE2. Mutated residues are shown as sticks, while contacting residues, including four residues on ACE2 and four tyrosine residues forming a hydrophobic cage are shown as lines. **(C)** Position of V1104L mutation and its role in conformational stabilization through cavity filling. **(D)** D339 and R346 are within the epitope region of class III antibody S309. Mutations on these two residues can contribute significantly to viral evasion. **(E)** Residues L455 and F456 (as sticks) surrounded by four tyrosine (as lines) are frequently targeted by RBD class I Mabs, such as CC12.1. In all panels, RBD is colored in green, ACE2 in brown, antibodies in magenta, cyan and yellow, and mutations are highlighted in red.

## DISCUSSION

The ongoing evolution of the SARS-CoV-2 spike protein continues to pose significant challenges to control the COVID-19 pandemic. Though BA.2.86 did not prove to have increased evasion of neutralizing antibodies in vaccinated and convalescent sera relative to prior variants^2–4,16,29^, subsequent variants, namely JN.1, have shown mounting escape^9,14,16,17^. Recent SARS-CoV-2 Omicron variants have been accumulating convergent mutations in the spike, particularly at key sites including L455, F456, and R346. These sites have critical roles in immunogenicity, receptor binding, and overall viral fitness^9,23,30,31^. Interestingly, despite its detrimental impact on ACE2 binding^14^, the single L455S mutation that differentiates JN.1 from BA.2.86 led to markedly more immune evasion^9,14,16,17^, sending the variant to dominance worldwide^12,19^. Here, we show that F456L (SLip) and R346T (FLiRT) contribute to further escape of JN.1-derived variants from neutralizing antibodies in all cohorts tested relative to their parental JN.1 (**Figures 2A-F**). FLiRT displayed the greatest decreases of nAb titer in bivalent immunized HCWs and in BA.2.86/JN.1 wave patients, likely due to the fact that the immunogens for both cohorts, i.e., WT, BA.4/5 or BA.2.86/JN.1, lack the R346T mutation (**Figures 2A-B**, **Figures 2E-F**). This notion is further supported by our finding that both FLiRT and KP.2 are neutralized better than SLip by the XBB.1.5-monovalent hamster serum samples, and that XBB.1.5 spike immunogen harbors the critical R346T mutation, which is present in both FLiRT and KP.2 but not SLip (**Figures 2C-D**). We and other groups have previously demonstrated that R346T is a critical site for monoclonal antibody binding and immune evasion^9,23^.

Another advantage of the R346T mutation is compensating for the loss of affinity for ACE2 caused by the L455S and F456L mutations (**Figure 6B**). Both the L455S and F456L mutations reduce hydrophobic contacts in the RBD, leading to a less binding for ACE2. R346T helps compensate for this reduction by strengthening conformational support in the receptor-binding motif (RBM) (**Figure 6A**). The consequences of these effects can be seen in the increased viral infectivity in FLiRT and KP.2 relative to SLip (**Figures 1C-D**). In particular, the KP.2 variant benefits from the effects of V1104L, which serves to further stabilize the spike conformation through hydrophobic internal cavity filling (**Figure 6C**), likely resulting in a less efficient transition from prefusion and postfusion spike, therefore impeding infectivity. We found that FLiRT and SLip were both resistant to neutralization by class III monoclonal antibody S309 like their parental variants BA.2.86 and JN.1. Accordingly, all of these variants including KP.2 possess the D339H mutation, which is situated directly in the center of the class III antibody epitope, creating a steric hindrance that abolishes S309 binding **(Figure 6D).**

The SLip variant displayed the lowest neutralizing antibody titers in the XBB.1.5-monovalent hamster cohort, likely due to the presence of the F456L mutation on top of its parental JN.1 containing L455S. The F456L mutation has occurred in several previous circulating variants, including FLiP, thus contributing to their strong immune evasion^32^. It is of note that this mutation is increasing in frequency among circulating variants^1^. F456L mediates immune evasion by altering key epitopes targeted by class I monoclonal antibodies **(Figure 6E)**, although other mutations (e.g., L455S) also contribute to this disruption of the class I epitopes. Notably, F456L is not present in XBB.1.5 spike immunogen, which could explain why SLip had the lowest neutralizing antibody titers in the XBB.1.5-monovalent hamster cohort. However, titers of XBB.1.5 monovalent hamster sera were still well above the limit of detection and only displayed a modest drop from JN.1, apart from SLip, suggesting that XBB.1.5 spike as an immunogen can still provide potentially effective protection against JN.1 -lineage variants. In addition, XBB.1.5-monovalent hamster sera exhibited less distance between the JN.1-lineage spikes, again suggesting that the XBB.1.5 spike as an immunogen could stimulate a broader antibody response than the WT+BA.4/5 bivalent vaccine (**Figures 4A-B**).

The BA.2.86/JN.1-wave cohort had similar antigenic mapping to the XBB.1.5 cohort, with even shorter distances between D614G and the JN.1 subvariants - and the JN.1 variants themselves were also closely clustered (**Figure 4C**). This pattern of response is similar to data presented by other groups, which showed that JN.1 infection, together with prior immunization, stimulates superior neutralizing antibody titers against JN.1 variants compared to BA.5, XBB, or XBB.1.5 breakthrough infections^9,14^. The shorter antigenic distance for our cohort, as well as the results from other groups, suggest that the JN.1 spike can and probably should serve as a more effective immunogen to stimulate neutralizing antibodies against JN.1-lineage variants. Overall, the difference between XBB.1.5 and BA.2.86/JN.1 spikes as immunogens against JN.1 variants (except SLip) appears negligible (**Figures 2C-F**, **Figures 4B-C**).

We did not find evidence of further enhanced nAb escape for KP.2 spike in three sets of sera relative to FLiRT and SLip (**Figures 2E-F**), which explains its increasing dominance in circulation. However, KP.2 acquired additional mutations in other regions of the SARS-CoV-2 genome^1^, including a T2283I mutation in non-structural protein 3 (nsp3) outside the papain-like protease (PLP) domain, which likely facilitates viral replication and/or modulate the host immune response^33,34^. Further investigations using authentic KP.2 and related JN.1 lineage variants shall help distinguish between possible mechanisms that endow KP.2 with a selective advantage over JN.1 and other subvariants in the pandemic.

Another interesting and somewhat surprising finding of this work is that most of the newly emerged JN.1 subvariants, especially SLip, exhibit decreased infectivity and cell-cell fusion activity in CaLu-3 cells compared to the parental JN.1 (**Figure 1**, **Figure 5**). Flow cytometric and western blotting analyses reveal decreased expression levels of the variant spike proteins on the plasma membrane of virus-producing cells, as well as reduced efficiency of spike processing by furin in the cell. Together, these findings could explain, in part, the observed infectivity and fusion phenotypes. Noticeably, these aspects of spike biology differ markedly from their parental BA.2.86 as well as some of the previously dominating Omicron variants, such as XBB.1.5 and EG.5.1, which exhibit increased spike processing, fusogenicity, and/or infectivity in CaLu-3 cells^35^. While the mechanism and implications of these differences in spike biology remain to be further investigated, our results suggest that some of these mutations, including F456L, R346T and V1140L, though beneficial for antibody escape, could negatively impact other aspects of spike biology, highlighting the critical tradeoff between immune evasion and viral fitness.

While the global COVID-19 pandemic has been declared over, SARS-CoV-2 continues to evolve and escape from host immunity elicited by vaccination and/or infections. Our data suggest that more recent JN.1 -lineage variants have altered properties in immune escape and biology, and they will continue to evolve. Our findings highlight the importance of continued tracking and characterization of emerging variants of SARS-CoV-2. Such studies are especially critical at this stage in the pandemic, when most people have been exposed to the virus at least once, if not several times, and thus have complex immunogenic backgrounds. Future vaccine development should consider JN.1 and/or closely related spikes as potential immunogen(s), though XBB.1.5 monovalent vaccines could still offer some protections.

### Limitations of Study

Our experiments make use of pseudotyped lentivirus bearing the spike protein of interest, not authentic infectious SARS-CoV-2 strains. However, the lentivirus model has proven to be an accurate reflection on neutralization of live SARS-CoV-2 and shown to instrumental for evaluating the efficacy of COVID-19 vaccines^36^. In addition, the timeliness of this work would not allow for thorough higher biosafety level three (BSL3) experiments to be conducted. The sample size of our cohorts, particularly the BA.2.86/JN.1 convalescents, is limited because of the IRB rules and restrictions. However, other similar studies have worked with cohorts of similar size^8,9,14^, and we have also published work with similar cohort sizes with reliable results in the past^5,16,21,23,37^. We recognize that homology modeling is not as precise as authentic cryoelectron microscopy (cryo-EM) structure, and the impact of key mutations on ACE2 interaction and antibody engagement would require confirmation by further structural studies. Nonetheless, data from these relatively small cohorts shall provide important insight into the biology of SARS-CoV-2 and offer timely guidance for future COVID-19 vaccine formulations.

## ACKNOWLEDGEMENTS

We wish to thank the Clinical Research Center and Center for Clinical Research Management of The Ohio State University Wexner Medical Center and The Ohio State University College of Medicine in Columbus, Ohio, especially Breona Edwards, Evan Long, J. Brandon Massengill, Francesca Madiai, Dina McGowan, and Trina Wemlinger, for collecting and processing the samples. We thank Tongqing Zhou at NIH’s Vaccine Research Center for providing the S309 monoclonal antibody. We thank Sarah Karow, Madison So, Preston So, Daniela Farkas, and Finny Johns in the clinical trials team of The Ohio State University for sample collection and other supports. In addition, we thank Moemen Eltobgy for assistance in sample processing. We thank Ashish R. Panchal, Soledad Fernandez, Mirela Anghelina, and Patrick Stevens for their assistance in providing the sample information of the first responders and their household contacts. We thank Peng Ru and Lauren Masters for sequencing and Xiaokang Pan for bioinformatic analysis. S.-L.L., D. J., R.J.G., L.J.S. and E.M.O. were supported by the National Cancer Institute of the NIH under award no. U54CA260582. The content is solely the responsibility of the authors and does not necessarily represent the official views of the National Institutes of Health. This work was also supported by a fund provided by an anonymous private donor to OSU. K.X. was supported by NIH grants U01 AI173348 and UH2 AI171611. M.C. was supported by an NIH T32 training grant (T32AI165391). J. L. was supported by NIH R01AI090060. J.S.B. was supported by award number grants UL1TR002733 and KL2TR002734 from the National Center for Advancing Translational Sciences. R.J.G. was additionally supported by the Robert J. Anthony Fund for Cardiovascular Research and the JB Cardiovascular Research Fund, and L.J.S. was partially supported by NIH R01 HD095881.

The authors have no competing interests to disclose.

S.-L.L. conceived and directed the project. R.J.G led the clinical study/experimental design and implementation. P.L. performed the experiments and data processing and analyses. K.X. performed molecular modeling and data analyses. D.J. led SARS-CoV-2 variant genotyping and DNA sequencing analyses. C.C., J.S.B., J.C.H., R.M., and R.J.G. provided clinical samples and related information. C.C.H, M.C., and J.L. provided hamster serum samples and associated information. P.L., J.N.F. and S.-L.L. wrote the paper. Y.-M.Z, L.J.S., E.M.O. provided insightful discussion and revision of the manuscript.

## STAR METHODS

### RESOURCE AVAILABILITY

#### Lead contact

Further information and requests for reagents and resources can be requested from the lead contact, Dr. Shan-Lu Liu.

#### Materials availability

Plasmids generated for this study can be made available upon request from the lead contact.

#### Data and code availability

This paper does not report original code. NT_50_ values and de-identified patient information will be shared by the lead contact upon request. Any other additional data can be provided for reanalysis if requested from the lead contact.

### EXPERIMENTAL MODEL AND SUBJECT DETAILS

#### Vaccinated and patient cohorts

The first cohort used in this study were healthcare workers (HCWs) at the Ohio State Wexner Medical Center that received at least 2 doses of monovalent WT mRNA vaccine and at least 1 dose of bivalent (WT+BA4/5) mRNA vaccine (n=10). Serum samples were collected under the approved IRB protocols 2020H0228, 2020H0527, and 2017H0292. All individuals received 2 homologous doses of monovalent mRNA vaccine, with 5 having received the Moderna formulation and 5 the Pfizer formulation. Four of these individuals received a homologous Moderna monovalent booster while 1 received a Pfizer monovalent booster. Four of the individuals in the Pfizer group received a homologous Pfizer monovalent booster while the last individual did not receive a monovalent booster. All individuals received 1 dose of bivalent mRNA vaccine encoding both WT and BA.4/5 spikes. Five received a Moderna bivalent dose and 5 received a Pfizer bivalent dose. The range of ages of individuals in this cohort was 27-46 years old with a median of 37. 5 males and 5 females were included. Blood was collected between 23-108 days post bivalent booster administration.

The next group used were golden Syrian hamsters (Envigo, Indianapolis, IN) vaccinated with monovalent XBB.1.5. The vaccine platform was a recombinant mumps virus expressing XBB.1.5 spike. The hamsters were vaccinated intranasally with 1.5 x 10^5^ PFU and administered a booster dose three weeks later. The hamsters were all 15 weeks in age. Blood was collected 2 weeks after administration of the booster dose.

The final cohort were individuals that were infected during the BA.2.86/JN.1 wave of infection in Columbus, OH (n=7). These sera samples were pulled from two sampling cohorts; the first were patients admitted to the ICU in the Ohio State University Wexner Medical center (n=3), the second were first responders and their household contacts part of the STOP-COVID cohort who were sampled when they became symptomatic (n=4). Samples were collected under the approved IRBs protocols 2020H0527, 2020H0531, 2020H0240, and 2020H0175. All samples were confirmed positive through RT-PCR and were collected between 11/23/2024 and 2/16/2024 which is when BA.2.86/JN.1 variants were dominant in Columbus, OH. Infecting variant was confirmed for a subset of samples through sequencing of nasopharyngeal swabs and next-generation sequencing using Artic v5.3.2 (IDT, Coralville, IA) and Artic v4.1 primers (Illumina, San Diego, CA).

See **Table S1** for full details on these cohorts.

#### Cell lines and maintenance

Cell lines used in this study include human epithelial kidney 293T cells (ATCC, RRID: CVCL_1926), 293T cells overexpressing human ACE2 (293T-ACE2) (BEI Resources, RRID: CVCL_A7UK), and human lung epithelial cell line CaLu-3 (ATCC, RRID:CVCL:0609). 293T and 293T-ACE2 cells were cultured in DMEM (Sigma Aldrich, Cat #11965-092) with 10% FBS (Thermo Fisher, Cat#F1051). and 0.5% penicillin/streptomycin (HyClone, Cat#SV30010). CaLu-3 cells were cultured in EMEM (ATCC, Cat 30-2003) supplemented the same way. For passaging, cells were first washed in PBS then detached with 0.05% trypsin+ 0.53mM EDTA (Corning, Cat#27106). All cells were cultured at 37°C in 5% CO_2_.

### METHOD DETAILS

#### Plasmids

All spike plasmids were in the pcDNA3.1 plasmid backbone with N- and/or C-terminal FLAG tags. The D614G plasmid was generated by GenScript Biotech via restriction enzyme cloning at Kpn I and BamH I sites and has a FLAG tag on both N- and C-termini. JN.1, FLip, SLip, FLiRT, and KP.2 were generated in-house through site-directed mutagenesis. The pNL4-3-intronGluc HIV vector was originally acquired from David Derse at NIH^38^.

#### Pseudotyped lentivirus production and infectivity

Pseudotyped lentivirus stocks were produced by transfecting 293T cells with pNL4-3 inGluc and the spike of interest. The transfection ratio was 2:1 vector to spike. Transporter 5 Transfection Reagent (Polysciences, Cat#26008-5) was used to carry out polyethyleneimine transfections. Pseudovirus was collected by taking the media off transfected cells 48 and 72 hours post-transfection. Equal volumes of media containing viral particles were used to infect target cells. The gLuc signals were measured by taking a portion of infected cell media and combining it with an equal volume of *Gaussia luciferase* substrate (0.1 M Tris pH 7.4, 0.3 M sodium ascorbate, 10 µM coelenterazine) and immediately reading luminescence in a Cytation 5 Imaging Reader (BioTek). 3 sequential readings were taken 48 and 72 hours post-infection and plotted. Relative infectivity was determined by setting the readout of D614G to 1.0.

#### Virus neutralization assay

The pseudotyped lentivirus neutralization assay was performed as described prior^36^. The infectivity of lentivirus stocks was pre-determined to ensure that similar amounts of infectious virus was used for each assay. Sera are serially diluted, initially diluting 1:40 followed by 4-fold dilutions (1:40, 1:160, 1:640, 1:2560, 1:10240) with one well left without sera. In the case of monoclonal antibody S309, the antibody was initially diluted to 12 µg/mL and serially diluted 4-fold with final concentrations 12, 3, 0.75, 0.19, 0.047 μg/mL. Equal volumes of normalized pseudovirus are then added to the diluted sera/antibody and incubated 1 hour at 37°C. The sera plus virus mixture is then used to infect 293T-ACE2 cells and luminescence readouts are taken 48 and 72 hours post infection. Neutralization titers at 50% are determined through least squares fit nonlinear regression using GraphPad v10 (San Diego, CA) normalized to the no sera/no antibody control.

#### Antigenic cartography analysis

Antigenic mapping was carried out using the Racmacs program v1.1.35 by following the workflow on the program’s associated GitHub entry (https://github.com/acorg/Racmacs/tree/master); Racmacs was run in R (Vienna, Austria). The user first inputs raw neutralization titers and the program log2 transforms them and creates a distance table representing antigenic distances between antigens (variant spikes) and individual sera samples. The program then uses this information to perform multidimensional scaling and plot the individual antigens (circles) and sera samples (squares) in two-dimensional space. The distance between points directly corresponds to fold changes in neutralization titers. One “antigenic distance unit” (AU) is equal to a two-fold change in neutralization titer and is represented by one side of the square. Optimization options in the program were kept default (2 dimensions, 500 optimizations, minimum column basis “none”). Maps were exported from the “view(map)” command and labeled using Microsoft Office PowerPoint.

#### Cell-cell fusion

293T cells are first transfected with plasmids of spike of interest and GFP. The cells are then detached 24 hours post-transfection and co-cultured with the target cell line. The cells are then co-cultured either 6.5 hours (293T-ACE2) or 4 hours (CaLu-3) and then fusion was imaged with a Leica DMi8 fluorescence microscope. Areas of fusion were quantified using the Leica X Applications Suite and by outlining edges of GFP signal. Scale bars represent 150µM.

#### Spike protein surface expression

293T cells used to produce pseudotyped lentivirus were collected using PBS plus 5mM EDTA to detach, and a portion of cells were fixed using 3.7% formaldehyde. Fixed cells were stained with anti-S1 polyclonal antibody (Sino Bio, T62-40591, RRID:AB_2893171) followed by anti-Rabbit-IgG FITC secondary (Sigma, F9887, RRID:AB_259816). Flow cytometry was run using an Attune NxT flow cytometer to determine surface expression of spike. Data was analyzed using FlowJo v10.8.1 software.

#### Spike protein processing

293T cells used to produce pseudotyped lentivirus were lysed using RIPA buffer (Sigma Aldrich, R0278) supplemented with protease inhibitor (Sigma, P8340). Samples were subjected to SDS-PAGE (10% polyacrylamide). Protein was transferred to a PVDF membrane then probed with anti-S2 (Sino Bio, T62-40590, RRID:AB_2857932) and anti-GAPDH (Proteintech, 10028230) antibodies. Secondary antibodies included anti-Rabbit-IgG-HRP (Sigma, Cat#A9169, RRID:AB_258434) and anti-Mouse-IgG-HRP (Sigma, Cat#A5728, RRID:AB_258232). Gels were imaged using Immobilon Crescendo Western HRP substrate (Millipore, WBLUR0500) on a GE Amersham Imager 600. Quantification of bands was determined using NIH ImageJ (Bethesda, MD).

#### Structural modeling and analyses

Structural modeling of the impact of spike mutations on ACE2 binding, conformational stability, and antibody evasion in the SLip, FLiRT, and KP.2 lineages was conducted using the SWISS-MODEL server. This analysis utilized published X-ray crystallography and cryo-EM structures (PDB: 7WK2, 8ASY, 7YAD, 6XC2) as templates. Key mutations were examined for their potential effects on these interactions, and the resulting models were visually presented using PyMOL.

### QUANTIFICATION AND STATISTICAL ANALYSIS

All statistical analyses described in the Figure legends were conducted using GraphPad Prism 10. NT_50_ values were calculated by least-squares fit non-linear regression. Error bars in (Figures 1C, 1D, 5C, 5D and 5F) represent means ± standard errors. Error bars in Figures 2A, 2C and 2E represent geometric means with 95% confidence intervals. Error bars in Figure 3A represent means ± standard deviation. Statistical significance was analyzed using log10 transformed NT_50_ values to better approximate normality (Figures 2A, 2C, 2), and multiple groups comparisons were made using a one-way ANOVA with Bonferroni post-test. Cell-cell fusion was quantified using the Leica X Applications Suite software (Figures 5A and 5B). S processing was quantified by NIH ImageJ (Figure 5G).

