## Supplemental Table for "Characteristics of JN.1-derived SARS-CoV-2 subvariants SLip, FLiRT, and KP.2 in neutralization escape, infectivity and membrane fusion"

### Table S1. Bivalent-vaccinated HCW and BA.2.86/JN.1-wave first responder cohorts

| | **Bivalent Health Care Workers** (n=10) | **BA.2.86/JN.1 Wave Patients** (n=7) |
|---|---|---|
| **Age in Years at Sample Collection [Median (Range)]** | 37 (27-46) | 51 (35-77) |
| **Gender** [n (% of Total)] | | |
| Male | 5 (50%) | 3 (42.9%) |
| Female | 5 (50%) | 4 (57.1%) |
| **Sample Collection Window** | Dec. 2022- Jan.2023 | Nov. 2023-Feb.2024 |
| **Vaccine status** [n (% of Total)] | | |
| 2-dose Moderna | NA | 2 (28.6%) |
| 3-dose Moderna | NA | 1 (14.3%) |
| 4-dose Moderna | NA | 1 (14.3%) |
| 1-dose Moderna +1-dose Pfizer bivalent | NA | 1 (14.3%) |
| 2-dose Pfizer +1-dose Pfizer bivalent | 1 (10%) | NA |
| 3-dose Pfizer +1-dose Moderna bivalent | NA | 1 (14.3%) |
| 3-dose Pfizer +1-dose Pfizer bivalent | 3 (30%) | NA |
| 3-dose Pfizer +1-dose Moderna | 1 (10%) | NA |
| 3-dose Moderna +1-dose Moderna bivalent | 4 (40%) | 1 (14.3%) |
| 2-dose Moderna +1 Pfizer +1-dose Pfizer bivalent | 1 (10%) | NA |
| Days from last vaccination | NA | 434 (34-892) |
| Days post the bivalent dose for recipients | 66 (23-108) | NA |
| **COVID-19 positive** [n (% of Total)] | 8 (80%) | 7 (100%) |
| Days before sample collection [(Median Range)] | 276.5 (182-994) | 7 (3-10) |
| **Infected Variants** | | |
| JN.1/BA.2.86 | NA | 2 (28.6%) |
| Undetermined | NA | 5 (71.4%) |

Summary of the demographic information for two cohorts used for neutralization experiments depicted in Figure 2. “NA” means the category is not applicable to the cohort.
